# ANALYSIS OF PHYTOEXTRACTS OF MEDICINAL PLANTS FOR ANTIMICROBIAL ACTIVITY AGAINST PATHOGENIC STRAINS OF MICROORGANISMS

**DOI:** 10.1101/2025.03.31.646455

**Authors:** Vasiliy A. Chokheli, Elena V. Butenko, Inna O. Pokudina, Irina M. Firsova, Anatoly S. Azarov, Mark O. Belyaev, Christina A. Tsymerskaia, Svetlana N. Sushkova

**Affiliations:** Southern Federal University (SFedU), st. Bolshaya Sadovaya, 105/42, Rostov-on-Don, Russia

**Keywords:** Artemísia absínthium, Thymus dimorphus, Salvia officinalis, Escherichia coli, Staphylococcus aureus, Klebsiella spp.

## Abstract

The study of aqueous and alcoholic phytoextracts of medicinal plants of the Rostov region: wormwood (*Artemisia absinthium*), thyme (*Thymus dimorphus*) and sage (*Salvia officinalis*) for pathogenic strains of *Staphylococcus aureus, Klebsiella sp*. and *Escherichia coli*. Various plant organs, such as flower, stem, leaf, and root, were used as phytoextracts. The antibacterial effect was provided by aqueous extracts from the leaves of *A. absinthium* in relation to *S. aureus*, as well as extracts of *S. officinalis* on *Klebsiella sp* strains. Alcohol extracts of these plants did not have an antibacterial effect on the studied strains of microorganisms.

## INTRODUCTION

Some biologically active substances of plants have antimicrobial properties, which makes it possible to use plant extracts and extracts against many human diseases. It is known that up to 70% of anticancer drugs are either completely of plant origin or contain components of plant origin. Currently, the use of plant biologically active substances is relevant, since many pathogenic microorganisms acquire high resistance to antibiotics. The decrease in the effectiveness of antimicrobial drugs and the emergence of antibiotic-resistant strains necessitated the search for new ways to combat persistent pathogens. In this regard, medicinal plants are of great interest, which can serve not only as a basis for the development of drugs with antimicrobial activity, but also act as a source of compounds with the necessary modifying activity against the persistent potential of the pathogen (Utkina, 2014).

It is known that in stressful living conditions for plants, the synthesis of substances of antioxidant activity is activated – low molecular weight peptides, organic acids, and flavonoid compounds. Thus, previous studies on the influence of various consequences of human economic activity, in particular, on the accumulation of flavonoids in wormwood grass harvested in urban and agrocenoses of the Central Chernozem region, showed strong variability in the results: with moderate anthropogenic load, there was an induction of the synthesis of flavonolic compounds, and with an increase in the content of flavonoids relative to other raw material samples in terms of on rutin, which can be explained by the inhibition of the plant’s antioxidant system (Dyakova et al., 2022).

In this regard, the purpose of this study was to analyze phytoextracts from various parts of medicinal plants for antimicrobial activity against pathogenic strains.

## MATERIALS AND METHODS

### Collection and preparation of plant raw materials for storage

The objects of the study were annual thyme plants (*Thymus dimorphus*), biennial wormwood plants (*Artemisia absinthium*), and two- and three-year-old sage plants (*Salvia officinalis*).

Collecting plant raw materials and preparing them for storage is an important initial stage in research. The plants were collected in the summer on the territory of the Botanical Garden of the SFedU.

The material was collected during the summer period: seeds were collected during their maturation, the aboveground parts of plants – during the period of maximum accumulation of BAS, starting from the flowering period until the formation of fruits. The collection was carried out in the daytime, in dry and sunny weather. They avoided collecting dusty plants. The collected plants were placed in containers. The collected vegetable raw materials were spread out on a clean cloth in a thin layer, it was removed from dead and damaged parts, from various impurities, dirt. The drying of plant raw materials was carried out in a ventilated room using natural heat with periodic stirring. After drying, the vegetable raw materials were packed in bags with labels with the name of the plant species, time and place of collection.

### Preparation of extracts

To prepare water extracts, fresh parts of the plant were crushed, 50 mL of distilled water was added to 5 g of vegetable raw materials and infused for 45 minutes at room temperature. The extracts were filtered through gauze and paper filters, then sterilizing filtration was performed using Filtropur S 0.45 filters (Sarstedt, Germany). Prior to performing microbiological studies, the infusions were stored at 6 ° C for no more than 24 hours.

Air-dried plants were used to prepare alcohol extracts. 50 mL of alcohol was added to 5 g of crushed plant parts and filtered through gauze and paper filters. The obtained extracts were stored in the refrigerator and used for microbiological studies for 30 days.

### Determination of antimicrobial properties of plant extracts

For microbiological studies, strains of microorganisms obtained from outpatient patients in Rostov-on-Don at the CJSC “Nauka” were used.

To determine the antimicrobial properties of plant extracts, a timing medium was poured into sterile Petri dishes at a temperature of 50-70° C, which were left under ultraviolet light for 15 minutes to achieve more accurate sterility. After the nutrient medium solidified, 100 µL suspensions of microorganisms were sown into Petri dishes using the continuous lawn method using a Drygalsky spatula. Antimicrobial activity was determined by diffusion into agar using filter discs. To do this, pre-sterile filter discs were placed in each Petri dish, on which the test solutions were applied, and the diameters of the microbial growth suppression zones were considered.

The antimicrobial properties of plant extracts were evaluated according to the following indicators:

1. The diameter of the microbial suppression zone is less than 3 mm – weak sensitivity to the substance.
2. The diameter of the microbial suppression zone is 4-10 mm – moderate sensitivity.
3. The diameter of the microbial suppression zone is more than 10 mm – high sensitivity.

Solutions of antibiotics were used for control. Antibiotics were used in accordance with the clinical recommendations for the relevant microorganisms.

## RESULTS AND DISCUSSION

### Determination of antibiotic resistance of the studied strains of microorganisms

To study the antibacterial properties of medicinal sage, wormwood, thyme representatives of gram-positive and gram-negative microorganisms were taken. *Staphylococcus aureus* is a gram-positive opportunistic bacterium of the genus *Staphylococcus*, which is the most common cause of staphylococcal infections, in particular nosocomial infections. Staphylococcus aureus can normally be located on the skin, nasal mucosa, and less often in the larynx, vagina, and intestines. They occur in 30% of healthy people.

*Klebsiella* is a genus of gram-negative encapsulated immobile bacteria belonging to the family *Enterobacteriaceae* of the order *Enterobacteriales*. The genus contains more than 12 species, of which *Klebsiella pneumoniae* and *K. oxytoca* are the most common. *Klebsiella* are part of the normal microbiota of the gastrointestinal tract, skin, and upper respiratory tract of healthy people. At the same time, they are one of the most common causes of both nosocomial and community-acquired infections, including urinary tract infections, bacteremia, pneumonia, neonatal abscess, and purulent liver abscess (Bardasheva, 2021).

*Escherichia* are polymorphic straight or slightly curved rods with rounded ends of medium size (length 2-6 microns and width 0.4-0.6 microns). The sticks are arranged singly, less often in pairs. They do not form a dispute. *Escherichia* are polymorphic straight or slightly curved rods with rounded ends of medium size (length 2-6 microns and width 0.4-0.6 microns). The sticks are arranged singly, less often in pairs. They do not form a dispute. The natural habitat of *Escherichia* is the distal intestine of humans, animals, birds, reptiles, and fish. *Escherichia* are sanitary- indicative microorganisms (Litusov, 2017).

The studied strains of microorganisms were tested for resistance to the main antibiotics. The results are presented in Table 1 and Figure 1.

**Table 1.** Results of testing of the bacterial strains studied for resistance to various.

| The microorganism | CLR | CD | LZ | CPM | LE | AMC |
| --- | --- | --- | --- | --- | --- | --- |
| *Staphylococcus aureus* gramm + | R | S | S | S | S | S |
|  | CIP | CTX | NIT | A/S | GEN | FO |
| *Klebsiella* gramm - | S | S | S | R | S | S |
| *E. coli* gramm - | S | S | S | R | S | S |

CLR – Clarithromycin; CD – Clindamycin; LZ – Linezolid; CPM – Cefepim; LE – Levofloxacin; AMC – Amoxiclav; CIP – ciprofloxacin; CTX – cefotaxime; NIT – nitrofurantoin; A/S – ampicillin/sulbactam; GEN – gentamicin; FO–fosfomycin.

**Figure 1.**
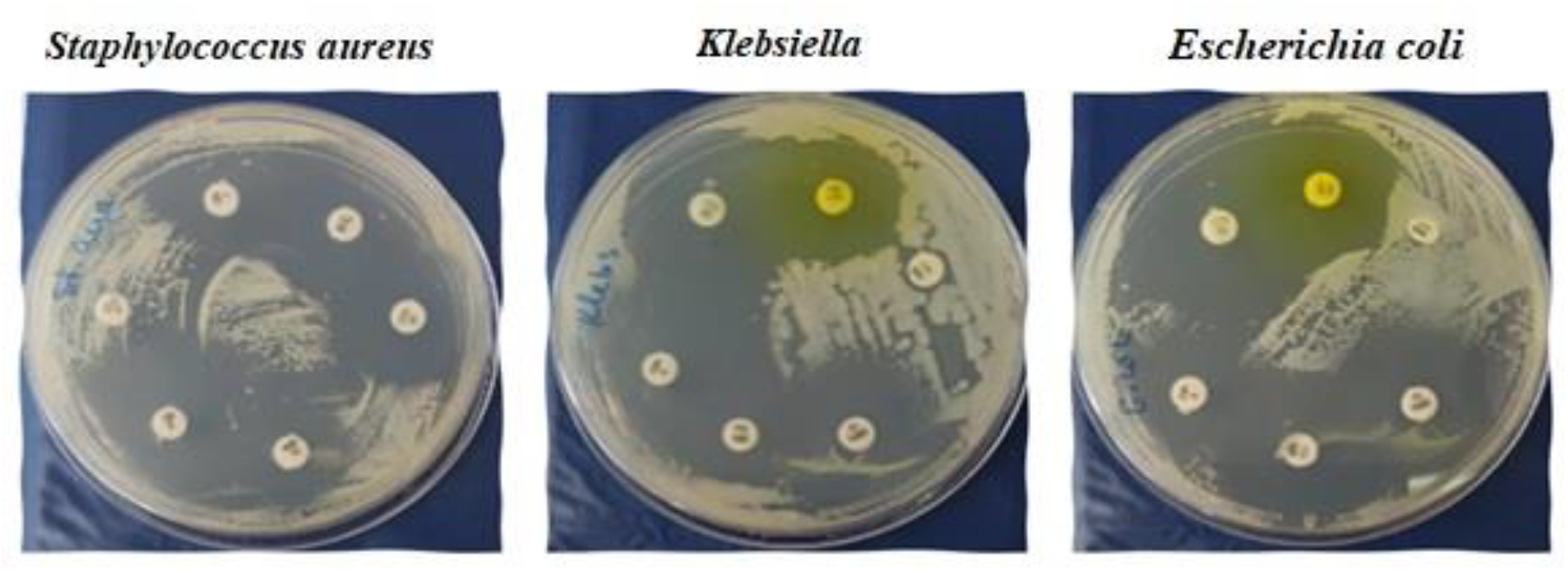
Results of testing of the provided strains for resistance to various antibiotics

### Determination of the antimicrobial activity of aqueous extracts

Different parts of the plant contain different secondary metabolites, and therefore their bactericidal properties may differ. As a result of the conducted research, it was found that aqueous extracts of stems, leaves, roots and flowers of thyme do not have antibacterial properties in relation to the studied microorganisms in the concentration used. The leaves of wormwood bitter in relation to Staphylococcus aureus had an antibacterial effect. The growth suppression zone was 4 mm. The results of the study of the antibacterial activity of an aqueous extract of an intact *Salvia officinalis* L., *Thymus dimorphus, Artemisia absinthium* plant on microorganisms of the species *Escherichia coli* and *Staphylococcus aureus* and the genus *Klebsiella* are presented in Table 2 and in Figures 2, 3.

**Table 2.** The results of the study of the antibacterial properties of alcohol extracts of *S. officinalis, T. dimorphus* and *A. absinthium*.

| Part of the plant | Growth suppression zone, mm |  |  |
| --- | --- | --- | --- |
|  | *S. aureus* | *E. coli* | *Klebsiella sp* |
| ***Artemisia absinthium*** |  |  |  |
| Flower | 0 | 0 | 0 |
| Stem | 0 | 0 | 1 |
| Leaf | 4 | 0 | 0 |
| Root | 0 | 0 | 0 |
| ***Thymus dimorphus*** |  |  |  |
| Flower | 0 | 0 | 0 |
| Stem | 0 | 0 | 0 |
| Leaf | 0 | 0 | 0 |
| Root | 0 | 0 | 0 |
| ***Salvia officinalis*** |  |  |  |
| 2-year-old plant |  |  |  |
| Flower | 0 | 0 | 0 |
| Stem | 0 | 0 | 0 |
| Leaf | 0 | 0 | 1 |
| Root | 0 | 0 | 0 |
| 3-year-old plant |  |  |  |
| Flower | 0 | 0 | 1 |
| Stem | 0 | 0 | 0 |
| Leaf | 0 | 0 | 0 |
| Root | 0 | 0 | 0 |

| Part of the plant | Growth suppression zone, mm |  |  |
| --- | --- | --- | --- |
|  | *S. aureus* | *E. coli* | *Klebsiella sp* |
| *Artemisia absinthium* |  |  |  |
| Flower | 0 | 0 | 0 |
| Stem | 0 | 0 | 0 |
| Leaf | 0 | 0 | 0 |
| Root | 0 | 0 | 0 |
| *Thymus dimorphus* |  |  |  |
| Flower | 0 | 0 | 0 |
| Stem | 0 | 0 | 0 |
| Leaf | 0 | 0 | 0 |
| Root | 0 | 0 | 0 |
| *Salvia officinalis* |  |  |  |
| 2-year-old plant |  |  |  |
| Flower | 0 | 0 | 0 |
| Stem | 0 | 0 | 0 |
| Leaf | 0 | 0 | 0 |
| Root | 0 | 0 | 0 |
| 3-year-old plant |  |  |  |
| Flower | 0 | 0 | 0 |
| Stem | 0 | 0 | 0 |
| Leaf | 0 | 0 | 0 |
| Root | 0 | 0 | 0 |

**Figure 2.**
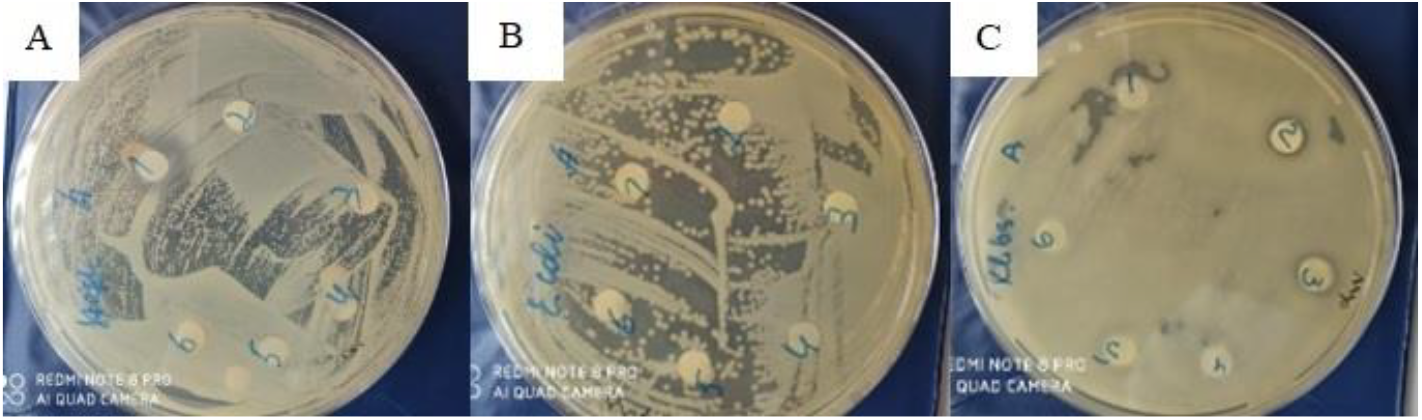
Antibacterial effect of wormwood extract (discs 1-2) A – *S. aureus*, B – *E. Coli*, C – *Klebsiella sp*.

**Figure 3.**
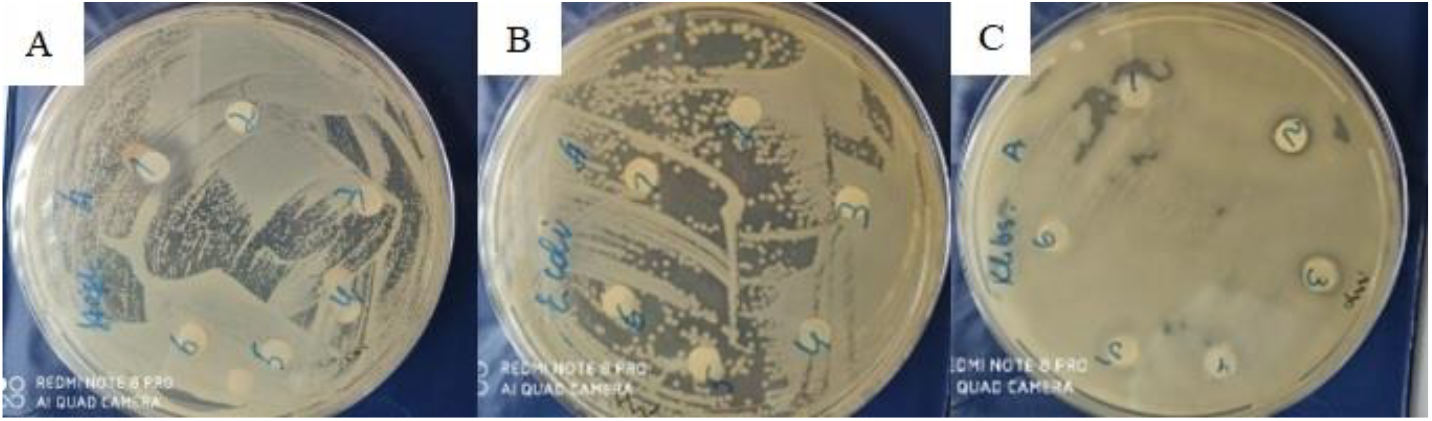
Antibacterial effect of sage extract (discs 2-6) A – *S. aureus*, B – *E. Coli*, C – *Klebsiella sp*.

As a result of the conducted research, it was found that aqueous extracts of stems, leaves, roots and flowers of medicinal sage do not have antibacterial properties in relation to the studied microorganisms in the concentration used. As can be seen from Table 2 and Figure 3, an aqueous extract of leaves of an intact medicinal sage plant of the 3rd year of life and an aqueous extract of flowers of the 2nd year of life in a concentration of 100% has weak antibacterial activity against Klebsiella spp. In other cases, no antibacterial activity is observed.

### Determination of antimicrobial activity of alcoholic extracts

The content of secondary metabolites in extracts also depends on the extraction method. Therefore, the antibacterial activity of alcoholic extracts of thyme, wormwood, and sage officinalis was also studied. As a result of the research, it was found that alcohol extracts of stems, leaves, roots and flowers of thyme and wormwood do not have antibacterial properties in relation to the studied microorganisms in the concentration used. The results are presented in table 3.

As a result of the conducted research, it was found that alcohol extracts of stems, leaves, roots and flowers of medicinal sage do not have antibacterial properties in relation to the studied microorganisms in the concentration used.

When comparing the bactericidal efficacy of aqueous and alcoholic plant extracts, it was revealed that it was the aqueous extract of wormwood leaves that had an antibacterial effect against staphylococcus aureus, while the alcoholic extract did not exhibit antimicrobial activity against all the microorganisms studied.

Two-year-old and three-year-old sage plants had the same antimicrobial activity, no differences were found between them.

Medicinal sage is one of the most used plants in traditional medicine. Studies of the antimicrobial potential of the *Salvia* genus have revealed wide variability depending on the sensitivity of microorganisms and the effectiveness of the tested compounds. Sage species rich in essential oils (such as *S. officinalis*) with volatile monoterpenoids as their main components have been reported to be effective antibacterial agents (Beheshti-Rouy et al., 2015). The essential oil of medicinal sage leaves contains a-thujone (27.66%), 1,8-cineol (8.07%), camphor (9.95%), viridiflorol (9.51%), and arabino-(4YU-methyl-glucurono)-xylan. 10 free and 11 bound amino acids have been identified in the liquid extract of medicinal sage leaves, of which tyrosine, serine, glutamic and aspartic acids are the dominant ones. The content of free amino acids is 0.48%, and the content of bound amino acids is 0.63% (Vovk et al., 2016). As a rule, Gram-positive bacteria are more sensitive to sage essential oil compared to other types of bacteria. The sensitivity of bacteria is related to the morphological structure and chemical composition of their shell (Horiuchi et al., 2007).

The chemical composition of *A. absinthium* L. varies depending on the growing area, plant part, ambient temperature, and plant age (Bordean et al., 2023). Many other researchers have found antibacterial activity in different species of Artemisia. In his study, Baykan Erel demonstrated a moderate effect of methanol extract of *A. absinthium* L. on *E. coli* (Erel et al., 2012). Sengul also reported antibacterial activity against E. coli for two types of extracts, aqueous and methanolic, from the aerial parts of *A. absinthium* L. (Sengul, 2011).

The genus *Thymus* is a very systematically complex group. We constantly have to deal with the existence of a large number of difficult-to-distinguish forms. Some authors identify many small species, while others suggest defining them into subspecies and hybridogenic forms. The characterization of species remains very complex due to the variability of not only chemical, but also morphological features of the genus species. The herb of the plant contains up to 0.6% essential oil, the main component of which is thymol (up to 42%). In addition, the essential oil contains carvacrol, n-cymol, α-terpineol, borneol. The studied components showed pronounced activity against a wide range of microorganisms – gram-negative (*Escherichia coli, Yersinia enterocolrtica, Pseudomonas aeruginosa, Salmonella choleraesuis, Salmonella typhimurium, Shigella dysenteria*) and gram-positive (*Listeria monocytogenes, Staphylococcus aureus, Staphylococcus epidermidis, Bacillus cereus, Enterococcus faecalis*), as well as molds (*Penicillium islandicum* and *Aspergillus flavus*, yeast *Candida albicans*). Along with these properties, thyme has been found to have a beneficial effect on the endoecological state of the gastrointestinal microflora because of controlling the growth of potential pathogens and stabilizing microbial eubiosis of the gastrointestinal tract (Vinokurova, 2016).

## CONCLUSION

As a result of the conducted research, it was found that alcohol extracts of stems, leaves, roots and flowers of medicinal sage do not have antibacterial properties in relation to the studied microorganisms in the concentration used.

When comparing the bactericidal efficacy of aqueous and alcoholic plant extracts, it was revealed that it was the aqueous extract of wormwood leaves that had an antibacterial effect against staphylococcus aureus, while the alcoholic extract did not exhibit antimicrobial activity against all the microorganisms studied.

Two-year-old and three-year-old sage plants had the same antimicrobial activity, no differences were found between them.

## Funding

The research was financially supported by a project “Molecular Biotechnology of Plants” within the framework of the Strategic Academic Leadership Program “Priority 2030” No. SP-12- 23-02.

## Notes

### Competing Interest Statement

The authors have declared no competing interest.

## REFERENCES

1. Bardasheva AV, Fomenko NV, Kalymbetova TV, Babkin IV, Chretien SO, Zhirakovskaya EV, Tikunova NV, Morozova VV. Genetic characterization of clinical Klebsiella isolates circulating in Novosibirsk // Vavilovskii Zhurnal Genet Selektsii. – 2021. – V 25(2). – P 234 – 245.

2. Beheshti-Rouy M, Azarsina M, Rezaie-Soufi L, Alikhani MY, Roshanaie G, Komaki S. The antibacterial effect of sage extract (Salvia officinalis) mouthwash against Streptococcus mutans in dental plaque: a randomized clinical trial // Iran J Microbiol. – 2015. – V 7(3). – P 173–7.

3. Bordean, M.-E.; Ungur, R.A.; Toc, D.A.; Borda, I.M.; Martis,, G.S.; Pop, C.R.; Filip, M.; Vlassa, M.; Nasui, B.A.; Pop, A.; et al. Antibacterial and Phytochemical Screening of Artemisia Species // Antioxidants. – 2023. - V 12. - P 596.

4. Dyakova N.A., Slivkin A.I., Korenskaya I.M. Features of Accumulation of Essential Oil in Bitter Wormwood Herbs of Flora of Voronezh Region of Russia // Drug development & registration. – 2022. – T 11(2). – C 140 – 144. 10.33380/2305-2066-2022-11-2-140-144.

5. Erel, S.B.; Reznicek, G.; Senol, S.G.; Yavasogulu, N.Ü.K.; Konyalioglu, S.; Zeybek, A.U. Antimicrobial and antioxidant properties of Artemisia L. species from western Anatolia // Turk. J. Biol. – 2012. – V 36. – P 75 – 84.

6. Horiuchi K, Shiota S, Hatano T, Yoshida T, Kuroda T, Tsuchiya T. Antimicrobial activity of oleanolic acid from Salvia officinalis and related compounds on vancomycin-resistant enterococci (VRE) // Biol Pharm Bull. – 2007. – V 30. – P 1147 – 1149.

7. Litusov N. V. CHastnaya bakteriologiya. Elektronnoe illyustrirovannoe uchebnoe izdanie. – Ekaterinburg: UGMU, 2017. – 707 s.

8. Sengul, M.; Ercisli, S.; Erzurum, T.; Yildizb, H.; Gungorc, N.; Kavaza, A.; Çetina, B. Antioxidant, Antimicrobial Activity and Total Phenolic Content within the Aerial Parts of Artemisia absinthum, Artemisia santonicum and Saponaria officinalis // Iran. J. Pharm. Res. – 2011. – V 10. – P 49 – 56.

9. Utkina T.M., Potekhina L.P., Kartashova O.L. The antimicrobial and antipersistent action of plant extracts from different species of Artemisia Southern Siberia // Siberian Medical Journal. - 2014. - №3. – P. 93–96.

10. Vinokurova O.A., Trineeva O.V., Slivkin A.I. Comparative characteristics of different types of thyme: the composition, properties and application (Review) // Development and registration of medicines. – 2016. – № 4(17). – p. 134 – 150.

11. Vovk G.V., Koshovyi O.M., Komissarenko A.M. The amino acid and monosaccharide composition of a dry extract from Salvia officinalis leaves obtained by complex processing // BіcниКфaрмaції. – 2016. – V 1 (85). - P 33 – 35.

